# *SituSeq*: An offline protocol for rapid and remote Nanopore amplicon sequence analysis

**DOI:** 10.1101/2022.10.18.512610

**Authors:** Jackie Zorz, Carmen Li, Anirban Chakraborty, Daniel Gittins, Taylor Surcon, Natasha Morrison, Robbie Bennett, Adam MacDonald, Casey R.J. Hubert

## Abstract

Microbiome analysis through 16S rRNA gene sequencing is a crucial tool for understanding the microbial ecology of any habitat or ecosystem. However, workflows require large equipment, stable internet, and extensive computing power such that most of the work is performed far away from sample collection in both space and time. Performing amplicon sequencing and analysis at sample collection would have positive implications in many instances including remote fieldwork and point-of-care medical diagnoses. Here we present *SituSeq*, an offline and portable workflow for the sequencing and analysis of 16S rRNA gene amplicons using the Nanopore MinION and a standard laptop computer. *SituSeq* was validated using the same environmental DNA to sequence Nanopore 16S rRNA gene amplicons, Illumina 16S rRNA gene amplicons, and Illumina metagenomes. Comparisons revealed consistent community composition, ecological trends, and sequence identity across platforms. Correlation between the abundance of phyla in Illumina and Nanopore data sets was high (Pearson’s r = 0.9), and over 70% of Illumina 16S rRNA gene sequences matched a Nanopore sequence with greater than 97% sequence identity. On board a research vessel on the open ocean, *SituSeq* was used to analyze amplicon sequences from deep sea sediments less than two hours after sequencing, and eight hours after sample collection. The rapidly available results informed decisions about subsequent sampling in near real-time while the offshore expedition was still underway. *SituSeq* is a portable and robust workflow that helps to bring the power of microbial genomics and diagnostics to many more researchers and situations.

## Introduction

Examining the microbiome of extreme and remote environments has increased our collective understanding of microbial physiology and diversity [1-2]. Collecting samples from these remote locations requires fieldwork that can be expensive and time consuming. Fieldwork is also logistically challenging as it can be complex to move people, equipment, and samples long distances, across borders, and through difficult terrain. In these situations, every sample taken is valuable in terms of the resources required for collection. Despite this, it is not always certain that the samples will address the research question, sometimes leaving researchers in situations where they are uninformed during sampling campaigns.

Sequencing of microbial genes (e.g., the 16S rRNA gene) is often used for environmental monitoring and medical diagnostics, as well as to identify indicators of environmental conditions, disturbances, and diseases [3-4]. Rapid analysis of 16S rRNA gene diversity could be used to quickly characterize a sample in circumstances when time or access to resources are limited. Sequencing with Oxford Nanopore technology (ONT), or “third generation” sequencing, is quickly gaining favour in the microbiological research community [5-8]. ONT allows for the continuous sequencing of long sequences of nucleotides, e.g., the full-length, 1500 bp 16S rRNA gene. While several ONT platforms have been developed and are in use, the MinION sequencer has gained recognition as relatively inexpensive and exceptionally portable. To exemplify its portability, the MinION sequencer has been used to sequence DNA in space [9-11], in remote field locations like the high Arctic [12] and during a ski touring expedition in Iceland [13]. In most previous cases of *in situ* sequencing, researchers have had to wait to analyze data until there was an internet connection or sufficient computing power available. This prevents the collection of meaningful information about field samples until much later, when field trips have ended and opportunities to adjust sampling strategies are gone. It would be advantageous to retrieve DNA sequence data in real time in the field, thereby informing decisions about whether to prioritize a site or move on to other sampling opportunities with the limited time and resources available, yet such methods are not currently available [5].

A drawback of ONT has been its low accuracy compared to conventional “second generation” sequencing-by-synthesis technologies like Illumina [14-15]. However, the accuracy of ONT is rapidly improving, with current estimates of >95% for raw read accuracy, leading to >99.99% consensus accuracies [16]. Furthermore, recent chemistry updates have enabled near-finished genomes without the need for polishing with short read Illumina sequences [17]. ONT is quickly approaching parity with other sequencing platforms in this regard [17-19].

Here, we present an amplicon sequencing workflow, *SituSeq*, designed to be used remotely with ONT. The workflow uses a MinION sequencer with a Flongle adapter, and a completely offline bioinformatics analysis pipeline with a pre-loaded database. The *SituSeq* method can be completed in less than an hour, and the entire process from DNA extraction to data visualization can be completed in less than eight hours. The workflow was tested during remote fieldwork in the NW Atlantic Ocean, assessing freshly collected deep sea sediments (>2000 m water column) ∼300 km offshore of Nova Scotia, Canada. Subsurface marine sediments contain a large amount of microbial biomass [20-21] yet harbour many uncultured and understudied taxa [22-24]. This is in large part due to the requirement for expensive resources (including ships for coring, personnel, and equipment), to collect samples in an often-limited time frame. These constraints make deep sea sediment sampling an ideal setting to test the implementation of this *in situ* sequencing method. Sequencing and analysis were carried out while at sea without internet connection and in turn informed subsequent sample collection during the remainder of the expedition. Refinement of the workflow was achieved using the same protocols back in the laboratory, and by comparing results to standard sequencing of amplicons and metagenomes from the same DNA using Illumina technologies. The code for running this analysis is available at https://github.com/jkzorz/SituSeq and in (S1 Data).

## Materials and Methods

Original DNA extraction, PCR, ONT library preparation, ONT sequencing, and subsequent data analysis were conducted at sea aboard the R/V *Atlantic Condor* in August 2021 [25]. This investigational effort resulted in the sequencing and analysis of four deep sea sediment *s*amples within hours of their retrieval, allowing the microbial community in each sample – including most notably the presence of hydrocarbon seep ‘indicator’ lineages [26] – to be assessed. Back in the laboratory, we then compared the results of the *SituSeq* ONT method conducted on a further 40 samples to the results of standard Illumina sequencing of 16S rRNA gene amplicons and metagenomes. Sequencing on all platforms was performed using the same extracted DNA. Below are the details of the *SituSeq* workflow and the comparison to Illumina sequencing.

### Sample collection and description

Forty marine sediment sub-samples were collected from different depth intervals within five push cores each approximately 30-40 cm in length (Fig 1). Push coring used a remotely operated vehicle (ROV; Helix Robotics), deployed from the R/V *Atlantic Condor*. The five push coring sites were labelled: “Purple Haze”, “Tiny Bubbles”, “Kilo”, “Clamshell”, and “175NW”. The first four showed visual evidence of hydrocarbon seepage and/or macroscopic fauna (Fig 1C), whereas “175NW” lacked distinguishable features similar to the majority of the abyssal sea floor (Fig 1B). Push cores were sectioned on board into 4 cm depth intervals, with sub-samples from each depth interval stored immediately at -80°C.

**Fig 1.**
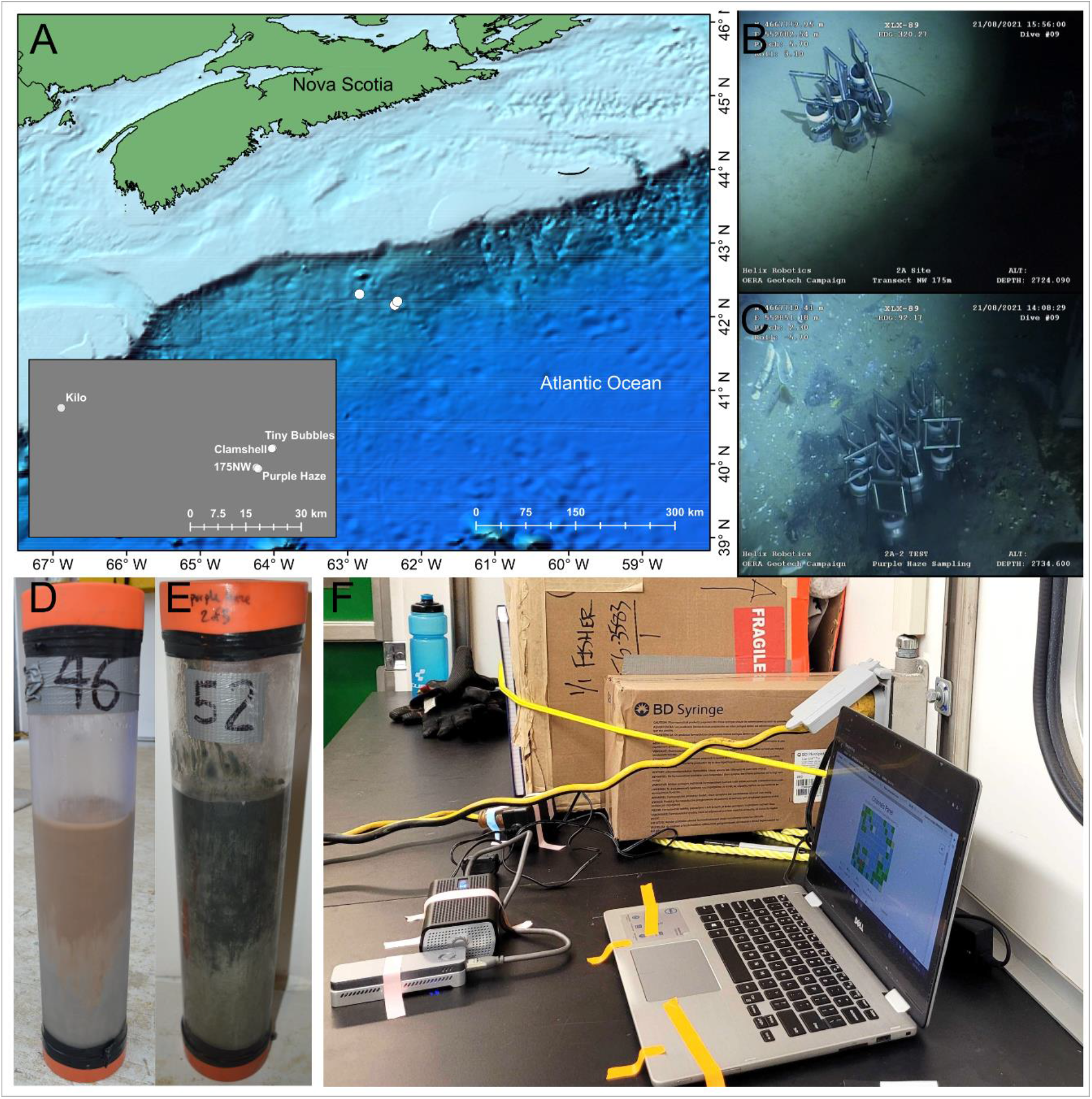
Research expedition on the Scotian slope offshore Nova Scotia. A) Sites sampled in the Northwest Atlantic Ocean. B) ROV footage from the background site “175 NW”. C) ROV footage from the hydrocarbon seep site “Purple Haze”. D) Image of the 31 cm long core taken from site “175 NW”. E) Image of the 36 cm long core taken from site “Purple Haze”. F) An ONT MinION sequencing run while at sea during the sampling expedition.

### Library preparation and sequencing methods

#### DNA extraction

DNA was extracted from the 40 marine sediment samples using the DNeasy PowerLyzer PowerSoil Kit (Qiagen, Germany) per the manufacturer’s instructions. Between 0.5-1 g of sediment was added to each lysis bead tube. Two 45 second rounds of bead beating using an Omni Bead Ruptor 24 bead beater (Omni-Inc, USA) at speed 5 were used to lyse cells. DNA was eluted in 70 µL of elution buffer (C6) following a two minute room temperature incubation. DNA concentration was measured with a Qubit fluorometric assay (ThermoFisher, USA).

#### Full length 16S rRNA gene barcoding PCR for Nanopore sequencing

Amplification of the full length 16S rRNA gene, clean up, and library preparation were performed using the 16S Barcoding Kit (SQK-RAB204, Oxford Nanopore Technologies, UK) per manufacturer’s instructions with minor modifications. This kit contains primers 27F/1492R for amplification of the full-length 16S rRNA gene (S1 Table), and has 12 barcoded primer pairs, allowing for the simultaneous sequencing of 12 samples. Instead of LongAmp Taq 2x master mix, KAPA HiFi HotStart master mix (Roche, Switzerland) was used. As per kit instructions, 10 ng of DNA per sample was used as template for the PCR except in samples with low extracted DNA concentrations (<1 ng/uL, determined using a Qubit fluorometer, ThermoFisher, USA) where at least 5 ng of template DNA was used. The PCR cycling conditions were altered slightly to accommodate the different polymerase enzyme and to improve extension conditions [27]. Alterations included longer denaturation and annealing phases (30s and 45s respectively in the cycles), and a higher temperature for the extension (increase from 65°C to 72°C). The thermocycler program used can be found in S2 Table.

A difficulty encountered during the barcoding and sequencing process was that barcode 8 and barcode 10 of the 16S Barcoding Kit (batch no. SE04.10.0020) consistently resulted in low-yield PCR products insufficient for downstream analysis. To remedy this, samples originally amplified with barcode 8 and barcode 10 were re-amplified with other barcodes to obtain enough material for the remainder of the protocol. Therefore, it is recommended that all barcoded primers are tested with positive controls prior to use.

PCR products were purified with AMPure XP beads (Beckman Coulter, USA), according to the instructions in the 16S Barcoding Kit. After PCR clean-up, the Qubit fluorometric assay was used to quantify DNA prior to pooling and normalizing libraries. Pooling of samples was done so that between 50-100 ng of total DNA was loaded in total, and between five and ten samples were included in each sequencing run. DNA was prepared for loading onto the Flongle adapter according to the ONT instructions. DNA prepared in this way from the 40 different sub-samples were run in batches that spanned six separate sequencing runs.

#### Nanopore sequencing

Sequencing was conducted using the MinION with a Flongle flow cell (R9.4.1) and with a MinIT (MNT-001) (Oxford Nanopore Technologies, UK) for basecalling. Sequences were locally basecalled using MinKNOW (v 4.3.20) (Oxford Nanopore Technologies, UK), connected to a Dell Inspiron 13-7378 laptop with 16 GB RAM and 512 GB SSD (Dell, USA). The length of the sequencing runs was variable and depended on Flongle flow cell quality and desired number of sequences per sample. In general, runs were continued until the active pores in the flow cells were depleted. To obtain 5,000 sequences per sample with six barcoded samples (30,000 sequences total), using a Flongle with around 50% of pores available (∼60 pores), approximately two hours of sequencing were required.

#### PCR amplification of V4 region of 16S rRNA gene and sequencing on an Illumina MiSeq

The same extracted DNA that was used for long-read ONT sequencing was used for short-read Illumina sequencing. Sample preparation and Illumina sequencing of the 40 samples was performed as previously described [28]. Briefly, the V4 region of the 16S rRNA gene was amplified using the 515F/806R universal primer set (S1 Table) [29-30]. The thermocycler programs used can be found in S2 Table. Amplicon samples were sequenced using Illumina’s v3 600-cycle (paired-end) reagent kit on an Illumina MiSeq benchtop sequencer (Illumina, USA).

#### Metagenome sequencing

To verify taxonomic community compositions, shotgun metagenomes were sequenced from nine out of the 40 samples using an Illumina NovaSeq (Illumina, USA). Libraries were prepared using a NEBNext Ultra II fragment library preparation kit (New England Biolabs, USA) with Covaris shearing (Covaris, USA). Libraries were then sequenced on a NovaSeq S4, 300 cycle run at the Centre for Health Genomics and Informatics (University of Calgary, Calgary, Canada), producing approximately 100M reads per sample.

### Data analysis

#### Nanopore analysis workflow

All code used for the analysis and instructions for analyzing data remotely and offline can be found at github.com/jkzorz/SituSeq. All software, packages, and databases used in the workflow require an internet connection to install, but once installed, can be run offline on a standard laptop (e.g., a Dell Inspiron 13-7378 laptop with 16 GB RAM and 512 GB SSD). An initial preprocessing script filters and trims reads, and then two analysis streams are offered: (1) taxonomic identification of all sequences with *dada2* [31] (program requirements: R), and (2) query of sequences against a pre-defined database of 16S rRNA gene sequences from species of interest using *BLASTn* (program requirements: R, *BLAST*). Both methods are described below (Fig 2).

**Fig 2.**
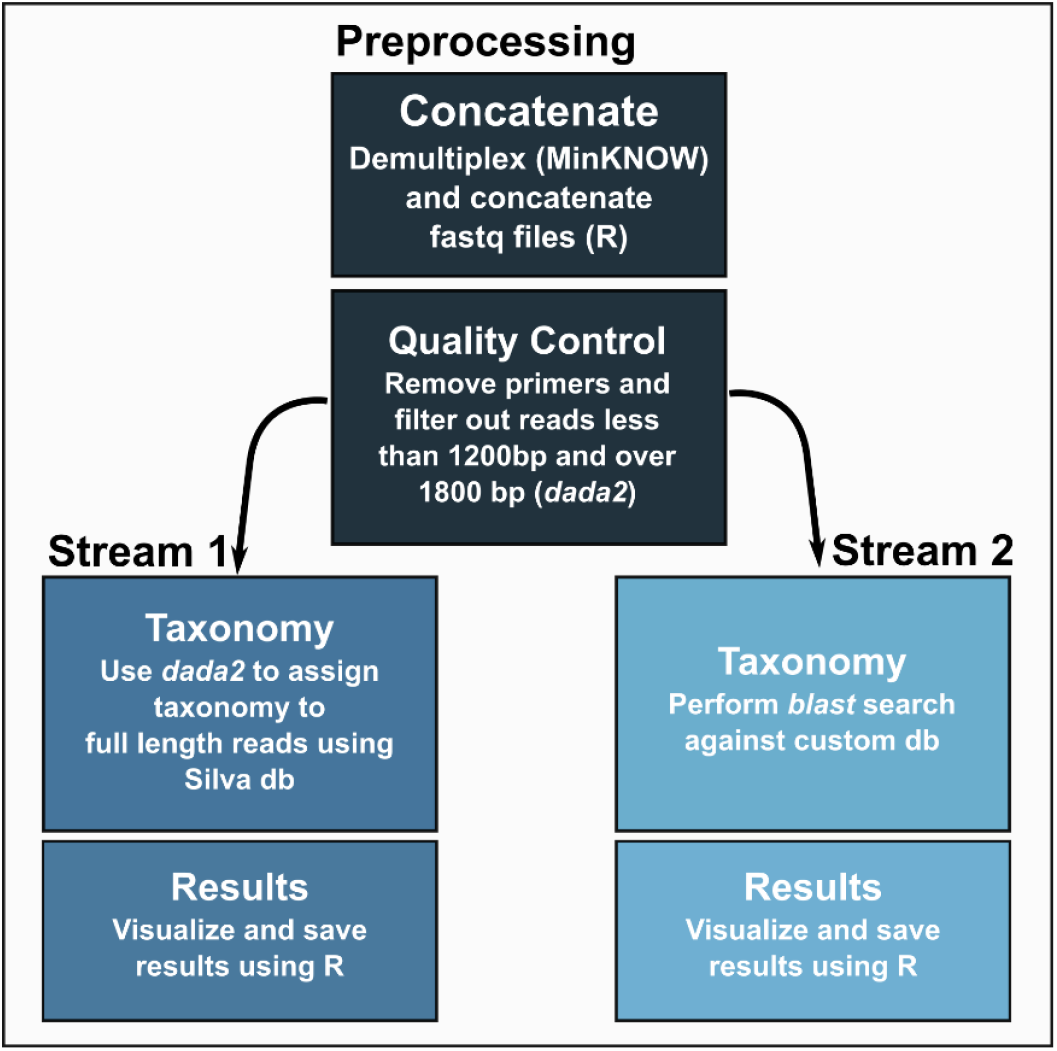
*SituSeq* bioinformatics workflow. The preprocessing script contains quality control steps that remove primers and filter sequences outside the specified length parameters. The Stream 1 workflow uses the *assignTaxonomy* function from *dada2* to assign taxonomy to all ONT sequences. The Stream 2 workflow performs a *BLASTn* search of the ONT sequences as queries against a custom database.

The preprocessing script concatenates separate sequence files from the same sample, and then uses the *filterAndTrim* command from *dada2* to remove primers and sequences longer or shorter than expected. For the present analysis, the first and last 100 bp of sequence were trimmed to remove primers and barcodes (trimLeft and trimRight), and then sequence reads were filtered using minLen = 1200 and maxLen = 1800 (filtering and trimming parameters are adjustable).

The first analysis method, Stream 1, is conducted entirely in R. It involves assigning taxonomy to full-length 16S rRNA gene sequences using a locally downloaded copy of the Silva 138.1 database [32] and the *assignTaxonomy* command from the program *dada2* (v1.20.0). Analyses were conducted on samples without subsampling sequences, or after subsampling to 1000 sequences.

The second method, Stream 2, involves a *BLASTn* identity search (v2.9.0+) of the sequences against a pre-defined database of 16S rRNA gene sequences belonging to species of interest for the project or field sampling in question. The 16S rRNA gene sequences included in the database in this instance belonged to hydrocarbon-associated bacteria identified in this study area [26] and in a previous study of cold seep sediments in the eastern Gulf of Mexico (S2 Data) [28]. The search database could include sequences from any number of species of interest to identify their presence in the samples being analyzed. The *BLASTn* command used required >97% identity, and the matches were filtered to remove any match with an e-value greater than 0.

#### Illumina analysis workflow

Samples sequenced on the Illumina MiSeq were analyzed using the *dada2* package in R [31] following the accompanying tutorial (https://benjjneb.github.io/dada2/tutorial.html). The samples were sequenced across two different MiSeq sequencing runs, such that the *learnErrors* and *dada* commands needed to be performed on each run separately. Two resulting ASV tables were then merged prior to taxonomic classification with the *mergeSequenceTables* command. Archaeal sequences were removed before comparing Illumina libraries with the ONT bacteria-specific full-length 16S rRNA gene libraries. The Silva 138.1 database was again used for taxonomic assignment, in the same manner as the ONT analysis. All code used in the analysis of the Illumina amplicon data is provided at github.com**/**jkzorz**/**SituSeq.

#### Reconstruction of 16S rRNA genes from metagenomes

Sequences underwent quality control using bbduk (BBTools suite; http://jgi.doe.gov/data-and-tools/bbtools), to remove the last base, adapters, contaminants, and low quality sequences. *PhyloFlash* v3.4 [33] was then used with the parameters: *-poscov -treemap -log -readlength 150*, to assemble and extract 16S rRNA sequences from the reads, and to assign taxonomy to those sequences using the Silva 138.1 database. Archaea and Eukaryote sequences were removed before calculating relative abundances of bacterial taxa. From the *phyloFlash* output, the files named LIBNAME.phyloFlash.NTUabundance.csv were used to calculate the relative abundance of taxa, and the files named LIBNAME.all.final.fasta, containing all assembled and reconstructed 16S rRNA gene sequences, were used for *BLAST* searches against ONT amplicon sequences.

#### Comparison of Illumina and Nanopore sequences

Taxonomic classifications and relative abundances of sequences were used to compare Illumina and ONT sequencing of the 16S rRNA gene from the same 40 samples. A three-way comparison between Illumina MiSeq amplicons, ONT amplicons, and *phyloFlash* sequences from Illumina metagenomes, was conducted for the nine samples that also had metagenomes. Many species found in deep sea sediments are poorly classified at finer taxonomic resolution, therefore the phylum level was used for these comparisons. The relative abundances of phyla identified using each sequencing technology were compared to assess any potential biases within the protocol. NMDS ordinations, ANOSIM tests, and Mantel tests were performed in R using the *vegan* package (v. 2.6-2) [34]. Differentially abundant phyla were identified based on a biserial correlation calculated using the *multipatt* function in the *indicspecies* package (v. 1.7.7) [35] in R. Differentially abundant phyla in different seabed locations were identified by grouping data separately for Illumina and ONT data sets. Combined ONT and Illumina data sets were used when identifying differentially abundant phyla based on sequencing technology. Pearson correlation between the relative abundance of phyla in the Illumina, ONT, and *phyloFlash* data sets was calculated using the *cor* function in R.

*BLAST* searches were done to directly compare sequence identities in a pair-wise manner for the three sequencing strategies (ONT 16S rRNA gene amplicons, Illumina 16S rRNA gene amplicons, and Illumina metagenomes). A custom searchable database was created from the ONT sequences using the command *makeblastdb*. Multiple *BLASTn* searches were also performed using the Illumina MiSeq and the *phyloFlash* metagenome sequences as queries. For the Illumina MiSeq searches, a requirement of 97% identity and a match longer than 230 bp was needed to be counted as a match. A second *BLASTn* search was done to search for 100% similarity between the Illumina MiSeq and ONT sequences. An unlimited (1000) amount of target sequence matches were included to allow for short read amplicons to match multiple ONT sequences. A *BLASTn* search with the parameters of 97% identity over 800 bp was used to query the reconstructed *phyloFlash* 16S rRNA gene sequences against the ONT sequence database.

## Results

### Illumina MiSeq 16S rRNA gene sequencing validation of Nanopore results

The *SituSeq* workflow presented here uses *dada2* in *R*, and optionally *BLAST* to analyze long-read ONT 16S rRNA gene amplicons offline and in near real-time. In order to validate the *SituSeq* workflow, the results of the *SituSeq* ONT 16S rRNA gene amplicon analysis were compared to the results from standard Illumina sequencing and analysis of 16S rRNA gene amplicons and metagenomes.

#### Community composition is similar with either amplicon sequencing method

Forty sediment samples spanning different depth intervals were collected from five push cores at sites on the Scotian Slope in the NW Atlantic Ocean (Fig 1). DNA extraction from sediment samples was performed in the same way both while at sea and in a land-based university laboratory, and 16S rRNA genes were sequenced using both long-read ONT and short-read Illumina sequencing technology. After filtering for length, amplicon libraries sequenced with ONT had an average of 18,630 reads (maximum: 38,121; minimum: 1,153). There were 745,171 full length ONT sequences retrieved in total (average length: 1403 bp after trimming and filtering). Illumina MiSeq amplicon sequencing resulted in a total of 912,570 bacterial sequences with an average of 22,814 bacterial reads per library (maximum: 84,462; minimum: 6,819), and an average length of 253 bp. In total, 7272 unique bacterial ASVs were formed from the Illumina reads.

A total of 66 phyla were identified in the ONT data set, and 65 phyla identified in the Illumina data set. The most abundant phyla on average across all 40 samples from the ONT data set were Proteobacteria (18.7%), Desulfobacterota (17.7%), Caldatribacteriota (11.5%), Campylobacterota (11.4%), and Bacteroidota (7.7%) (Fig 3A). Illumina results for the same samples grouped at the phylum level were similar with the most abundant being Proteobacteria (26.0%), Desulfobacterota (16.7%), Caldatribacteriota (12.1%), Planctomycetota (6.9%), and Bacteroidota (6.1%) (Fig 3B). Within the ONT data set, 2.9 ±1.2% of the community had no taxonomic classification at the phylum level, compared to 0.8% ± 0.7% of the community within the Illumina data set. However, at the genus level, there were fewer sequences without classification within the long-read ONT data set (65% ±12% of sequences and relative abundance) than in the short-read Illumina data set (80% of ASV sequences, collectively comprising 65% ± 11% relative abundance of the Illumina data set).

**Fig 3.**
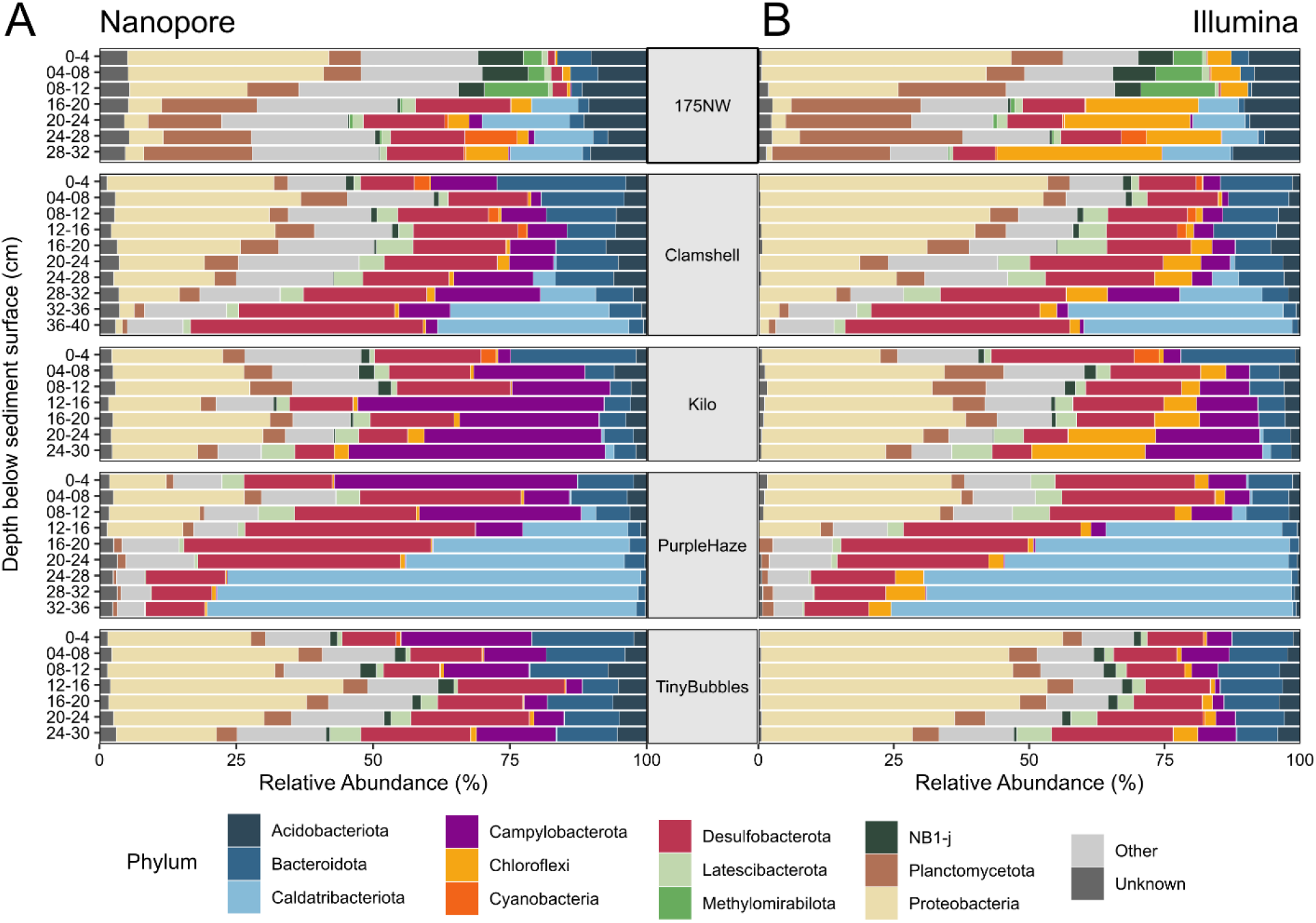
Relative abundance of most abundant phyla. A) Abundance of phyla across samples from ONT data set. B) Abundance of phyla across samples from Illumina data set. Unknown represents the sequences not identified at the phylum level and other represents the less abundant phyla.

Overall, there was high correlation between the relative abundance of phyla from ONT and Illumina data sets (Pearson’s r = 0.905) (Figure 4A). There were, however, differences between the sequencing technologies in terms of the relative abundance of certain phyla. Chloroflexi had a much higher relative abundance (6.4x ± 4.2x higher) in samples sequenced with Illumina technology than in samples sequenced with ONT (Fig 4B). The phylum Campylobacterota, in contrast, was more abundant in ONT samples (2.8x ± 1.8x) compared to Illumina samples. S3 Table contains the phyla that were differentially abundant between the sequencing methods.

**Fig 4.**
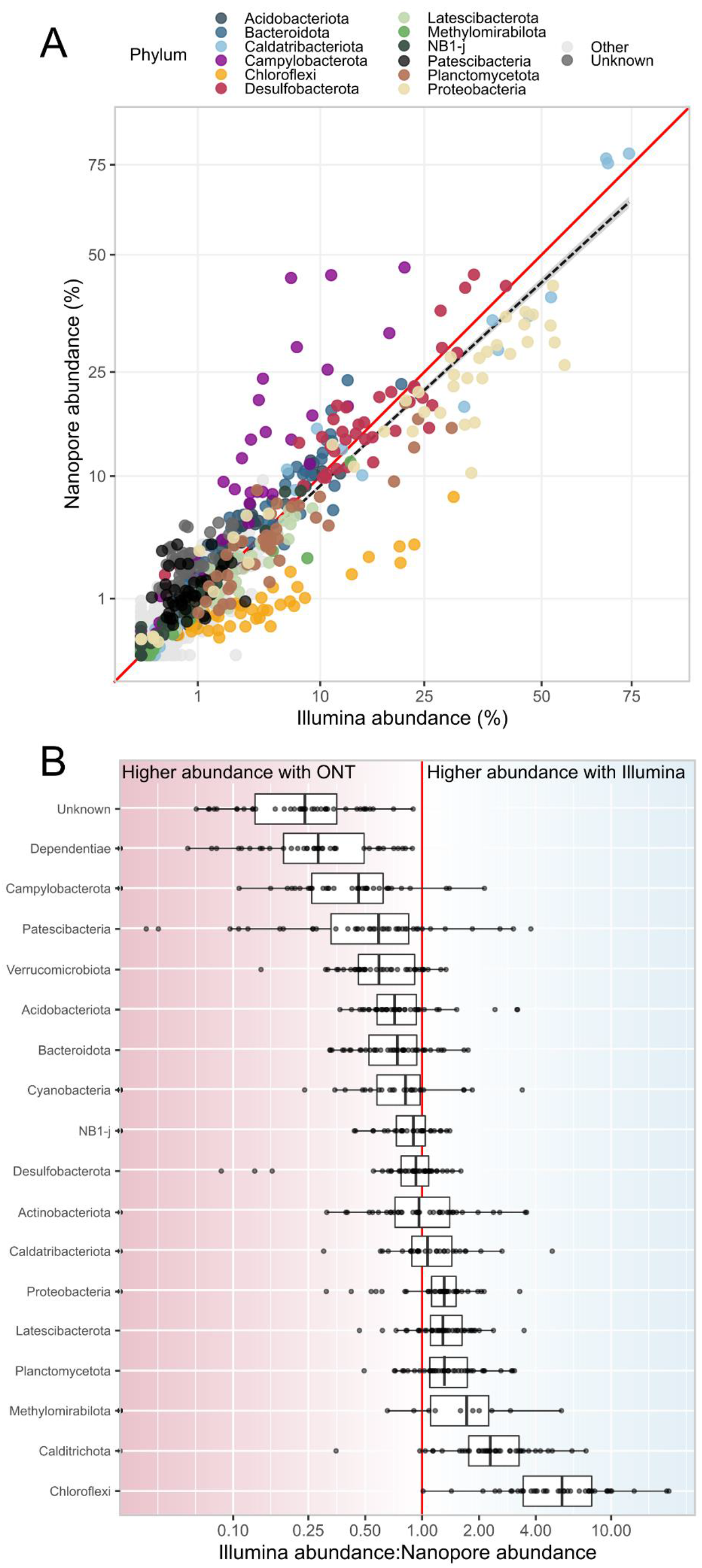
Comparison of ONT and Illumina amplicon sequencing. A) Correlation between abundance of phyla sequenced with ONT (y-axis), and Illumina (x-axis) technology (Pearson’s r = 0.905). Abundant phyla are coloured, unknown represents the sequences not identified at the phylum level, and other represents the less abundant phyla. The black dashed line shows the linear relationship between ONT and Illumina abundances, and the red line shows a 1:1 ratio. The axes have been square root transformed. B) The ratio of Illumina abundance to ONT abundance of select phyla. Ratios for individual samples are overlaid on the boxplots. The red line shows a 1:1 ratio.

#### Beta diversity is similar with either amplicon sequencing method

ONT and Illumina amplicon data sets were combined at the phylum level to determine the effects of sequencing technology (a combination of library preparation and sequencing platform) on broad ecological conclusions (Fig 5). Sequencing technology had a significant effect on microbial community composition (ANOSIM p = 0.008), but the strength of this impact was small (ANOSIM statistic = 0.06) and did not mask the effect of sampling location (site) (ANOSIM statistic = 0.38, p < 1e^−4^). Differences in microbial communities between samples in the combined data set were significantly correlated with differences between sediment depth intervals in the seabed (Mantel test: 0.33, p < 1e^−4^). When ONT and Illumina data sets were evaluated separately, the effect size of location and correlation with depth were significant (p < 1e^−4^) and similar to the combined data sets. ANOSIM statistics of location for ONT-only and Illumina-only data sets were 0.36 and 0.37, respectively, and Mantel statistics for ONT-only and Illumina-only data sets were 0.30 and 0.35, respectively. Therefore, the major ecological trends in the data were still evident despite differences in primers and sequencing platforms.

**Fig 5.**
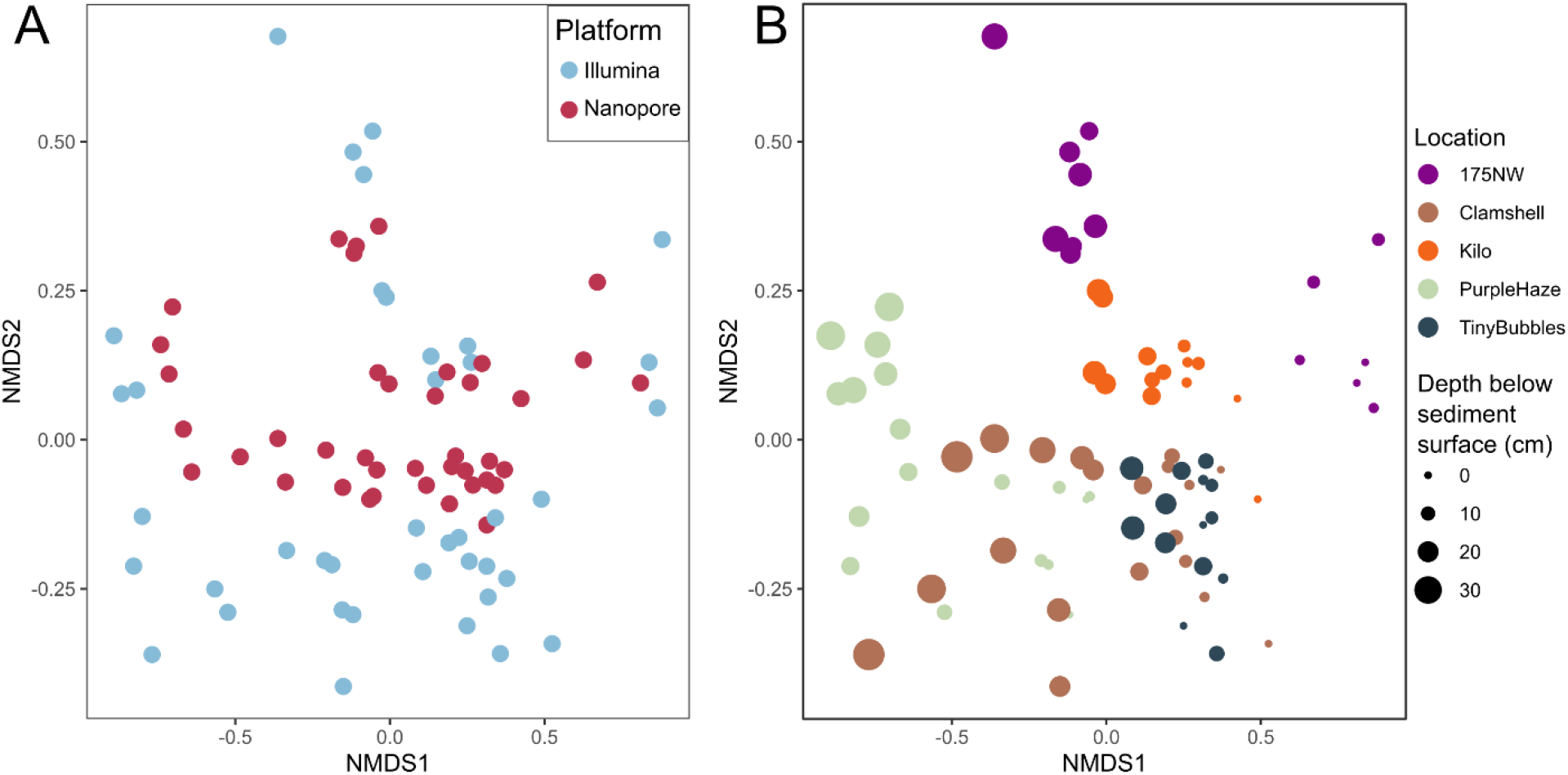
NMDS plots with combined ONT and Illumina data sets. The same NMDS plots are shown with samples (points) coloured and sized based on different parameters. A) Sample colour indicates sequencing technology. B) Samples coloured on location, with size proportional to depth below the sediment surface. Stress = 0.0999.

#### Sequence identities are high between amplicon sequencing methods

To compare ONT and Illumina data sets at a finer taxonomic resolution, a *BLAST* search was performed using the Illumina ASV sequences as queries against a custom database composed of the ONT sequences. This revealed that 5,303 out of the 7,272 Illumina bacterial ASVs (73%) had *BLAST* hits with greater than 97% percent identity among the ONT sequences. Of the top 1,000 most abundant Illumina ASVs (comprising 83% of the Illumina data set relative abundance), only 20 ASVs did not match to any of the ONT full length sequences. This indicates that the most abundant species in the community were identified the ONT data set with high sequence agreement.

A *BLAST* search identifying sequence matches with 100% identity over 230 bp Illumina sequences was also conducted to evaluate perfect matches. Of the 7,272 Illumina ASVs, 567 (7.8%) had *BLAST* hits with 100% identity to a full length ONT sequence. Some of these ASVs had 100% identity to multiple ONT full length sequences (i.e., owing to ONT sequences differing from each other in other areas of the 16S rRNA gene), such that 1,249 of the full-length ONT sequences matched perfectly to ASVs from the Illumina data set. Of the top 21 most abundant Illumina ASVs, 20 had 100% identity matches to ONT sequences (the exception being ASV8 from Chloroflexi), and in general, the more abundant Illumina ASVs had a higher number of 100% identical hits to the ONT sequences. For example, ASV1 (Caldatribacteriota) and ASV2 (*Sulfurovum*), had 100% matches to 87 and 37 unique ONT sequences respectively. This highlights the range of sequence diversity that is missed when using shorter variable regions and shows the potential of full length 16S rRNA gene sequencing for greater taxonomic resolution past what is possible with short-read amplicons.

### Shotgun metagenome taxonomy validates Nanopore results

To assess how amplicon-based methods compared to primer-free methods, 16S rRNA genes were reconstructed from nine Illumina metagenomes and were compared to the ONT and Illumina amplicon sequences. At the phylum level, the Illumina metagenome relative abundances were highly correlated with the ONT amplicon relative abundances (Pearson’s r = 0.876). Similar to the ONT-Illumina amplicon comparison, the phylum Chloroflexi was much more abundant in the Illumina metagenome data set, and the phylum Campylobacterota was more abundant in the ONT data set (Fig 6), suggesting that the ONT primers for 16S rRNA genes under- and over-represent these two phyla, respectively. The phylum Patescibacteria was much higher in the Illumina metagenome data set than in either ONT or Illumina amplicon libraries (Fig 6), suggesting that both sets of PCR primers may result in underestimation of this phylum in amplicon libraries [36]. Similarly, Actinobacterota, Firmicutes, and Poribacteria [37] were significantly more abundant in the Illumina metagenome data set than in both ONT and Illumina amplicon libraries (Fig 6).

**Fig 6.**
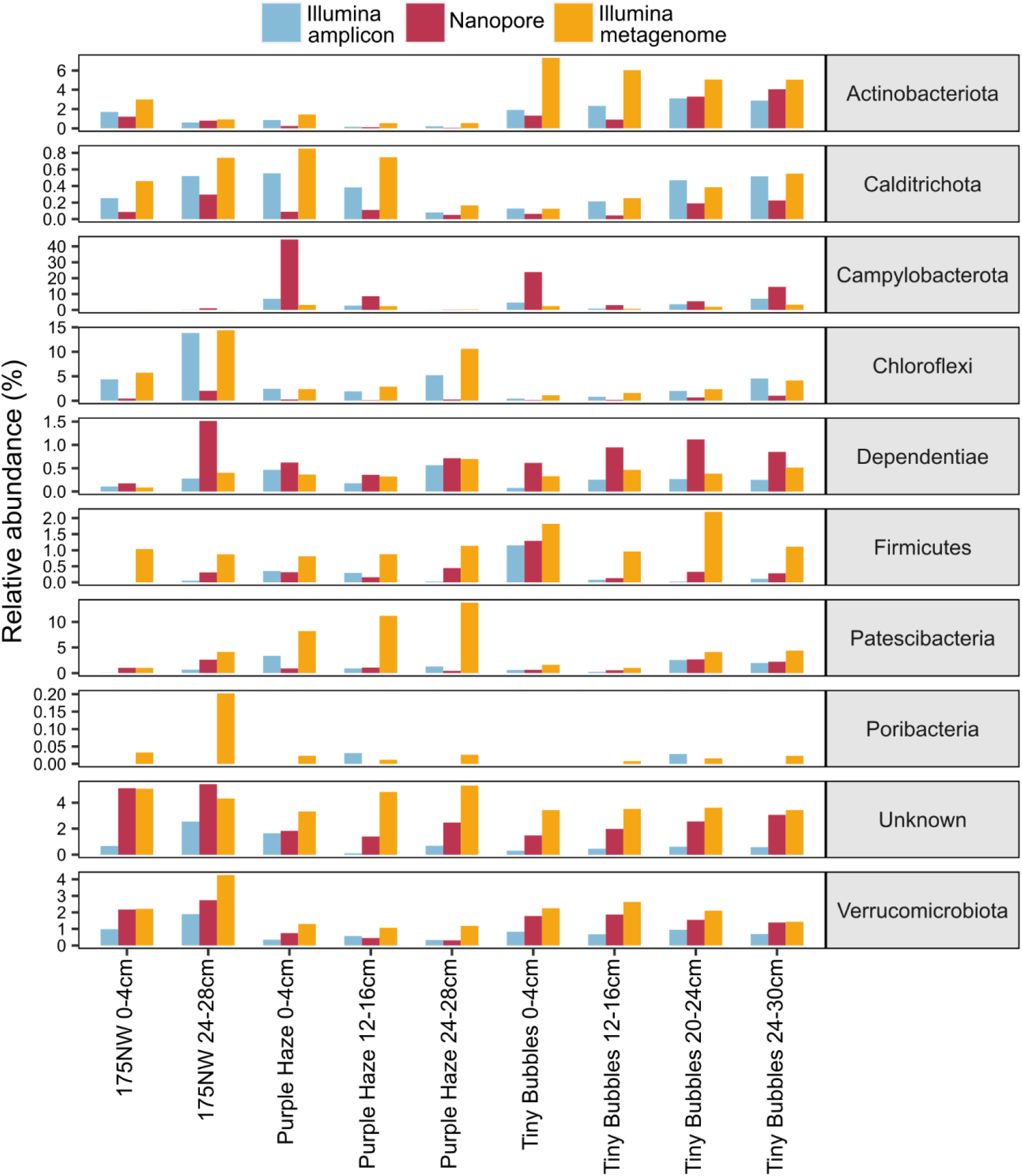
Comparison of relative abundance of a selection of differentially abundant phyla between ONT amplicon, Illumina amplicon, and Illumina metagenome data sets. Note the different scales of the y-axes.

A *BLAST* search was performed using the 16S rRNA gene sequences reconstructed from the metagenomes as queries against a database made from the ONT amplicon gene sequences. Of the 407 16S rRNA sequences assembled from the metagenomes that were over 800 bp long, 284 (69.8%) had *BLAST* hits with greater than 97% identity to the ONT sequences. This shows that the majority of ONT sequences, even without error correction, would fall within the conventional lowest taxonomic unit cluster, the OTU (operational taxonomic unit), with near full-length Illumina 16S rRNA gene sequences.

### User defined database for targeted sequence identification without an internet connection

In addition to assigning taxonomy to all ONT reads based on Silva version 138.1 (Fig 3A), the *SituSeq* workflow presented here supports an additional option (Stream 2) to use *BLAST* to match ONT amplicon sequences to a pre-populated user-defined database containing sequences of interest. This enables meaningful context-specific data analysis without an internet connection. One of the goals of the research expedition was to identify sediment samples in close proximity to hydrocarbon seepage using the presence of bacterial taxa previously found to be associated with hydrocarbon seeps. Thus, in our case, a database of 21 hydrocarbon-associated bacterial lineages from deep sea cold seeps was used [28]. All 21 of these sequences had *BLAST* hits to ONT sequences with >97% identity. Hits were found from a total of 4541 ONT sequences in 32 samples, with the sites 175NW and Purple Haze, having the highest average relative abundance of hydrocarbon-associated lineages (2.8%) (Fig 7). Hits at the 175NW site were only found ≥16 cm depth, and the relative abundance of the hydrocarbon-associated lineages in the Purple Haze site increased greatly ≥12 cm depth. The Kilo site also showed high abundance of hydrocarbon-associated species (2.3% on average), while Clamshell site had a higher abundance of hydrocarbon-associated species in deeper samples. Tiny Bubbles site had the fewest hits with only 21 matches across all depths. In this particular case, the Kilo site, as well as deeper samples from 175NW, Purple Haze, and Clamshell sites, were deemed to be of interest for further investigation of hydrocarbon-associated species during the field expedition, whereas the Tiny Bubbles site was de-prioritized.

**Fig 7.**
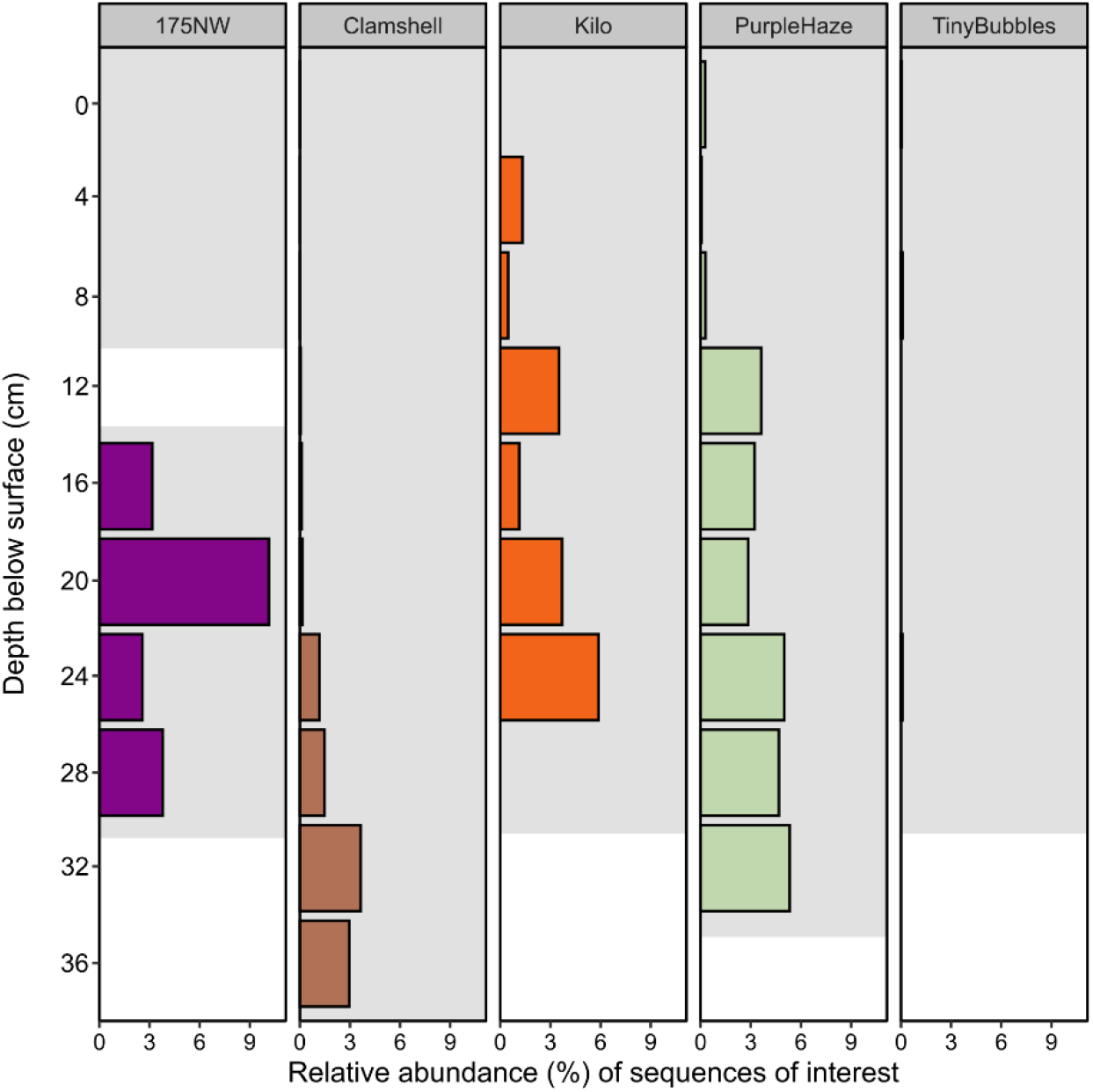
Combined relative abundance of 21 taxa of interest in each sample. The shaded grey area shows the depths of the samples that were sequenced for each core.

### Subsampling of Nanopore amplicon reads to reduce the time required for the overall workflow

Taxonomic assignment of 16S rRNA gene sequences (Stream 1) is the computational bottleneck in the *SituSeq* analysis workflow. To speed this up and reduce the amount of time needed for sequencing, a smaller subsampled data set was assessed to see whether fewer sequences would increase computational efficiency without affecting conclusions about microbial community structure and ecology. ONT libraries were subsampled to 1,000 sequences and community composition based on this was compared to the community composition based on the non-subsampled data (average library size of 18,629 sequences).

Ecological trends derived from the 1,000 read sequencing depth were very similar to those observed with non-subsampled larger libraries (S4 Table). Across all 40 libraries, 61 phyla were identified after subsampling. This is only 5 fewer than the 66 phyla detected in the larger data set, despite the removal of 705,171 reads (95% of the ONT data set). The effect of location (site) on community structure was still significant (ANOSIM stat = 0.382, p < 1e^−4^), as was the relationship between depth and community structure (Mantel test: 0.333, p < 1e^−4^). The same phyla were picked out as differentially abundant between locations (S4 Table). In our case, as little as 1000 sequences per sample adequately captured the ecological trends in our data set.

## Discussion

The ability to rapidly sequence and analyze samples completely offline without an internet connection, offers major advantages in settings such as remote field work and rapid point-of-care diagnostics. This study demonstrates robust offline analysis, through the *SituSeq* workflow, of 16S rRNA gene sequences obtained using Oxford Nanopore sequencing and the highly portable MinION sequencer. The method and interpretations were verified here by comparing *SituSeq* results to standard Illumina MiSeq sequencing of the V4 region of the 16S rRNA gene, and to Illumina NovaSeq metagenome-derived 16S rRNA gene sequences. Overall, there was very good agreement between the methods, with the main discrepancies likely due to PCR primer selection rather than the sequencing platform [38-40]. *SituSeq* is designed to be simple enough to be implemented by users with little bioinformatics experience. The simplicity of the workflow is beneficial for remote deployment where teams of experts can be small, yet important decisions must be made rapidly and accurately.

Several workflows with varying strategies and goals currently exist for the analysis of ONT-sequenced 16S rRNA gene data. For example, the *spaghetti* pipeline [41] designed to aid targeted bioprospecting in the field - a similar objective to the present study. The *spaghetti* pipeline comprises multiple steps including removal of primers and adapters with *Porechop* (no longer supported), filtering with *Nanofilt* [42], quality control with *Nanostat* [42], and *minimap2* [43] for mapping long reads to the Silva database. Thus, the *spaghetti* workflow requires installation of multiple separate programs, and uses *minimap2* for taxonomic assignment, increasing the computing power and bioinformatics expertise required for execution [43]. Recently, Curry et al. (2022) [44] developed *Emu*, a command-line workflow for community profiling of 16S rRNA gene ONT sequencing data. The method relies on an expectation-maximization algorithm to correct for the inherent sequencing errors of ONT and uses *minimap2* to map long reads to a database. *Emu* produced highly accurate results compared to conventional sequencing methods but was computationally intensive, requiring more threads and RAM than is usually available on a standard fieldwork laptop to analyze diverse communities. EPI2ME is the standard 16S rRNA analysis and annotation software from Oxford Nanopore and is accessed through a graphical user interface (https://epi2me.nanoporetech.com/). However, it is cloud based and requires an internet connection to use. Due to the ease of offline use and minimal requirements for software installation and computing power, *SituSeq* is a valuable addition to this suite of Nanopore 16S rRNA gene analysis workflows.

In the present study, the uncultured and unclassified nature of many of the resident microorganisms was overcome by using a customized database of indicator sequences derived from other deep sea cold seep sites [26,28]. In other situations, well-designed local databases queried from a standard laptop could be an important strategy for rapid identification of environmental or medical species of interest. Consider the context of rapid diagnostics, which currently utilizes rapid isothermal PCR reactions without sequencing but rather employing primers that are specific for a given bacterial pathogen [45-46]. In the case of a negative result, those assays would need to be repeated with different primers targeting other specific pathogens. Application of *SituSeq* with a well-designed database of potential pathogenic candidates could rapidly uncover and diagnose mysterious cases where access to, for example, a large research hospital is not feasible [47].

The ability to characterize a microbiome in real-time could greatly aid many fieldwork expeditions and help researchers make informed decisions about which samples to focus on. For instance, real-time results identifying taxa of interest would aid in the selection of samples for more in-depth analysis requiring more material (e.g., metagenomics, metaproteomics). Studies requiring enrichment or incubation from environmental samples would also benefit from knowing the contents of the inoculant beforehand, and methods like *SituSeq* could be used to target samples containing coveted species for cultivation [41]. In addition, the ability to sequence at source, could potentially reduce the number of samples needing to be stored and transported (reducing the cost and risks associated with these tasks), and could aid in the characterization of sensitive samples [48].

The *SituSeq* workflow presented here is designed to produce rapid results that can be accessed in the field. The workflow results are highly comparable to what one would receive from a conventional 16S rRNA gene amplicon analysis of a variable region using second generation sequencing technologies. Portable ONT sequencing in combination with easy and reliable workflows, like the one presented here, will expand the accessibility of sequencing beyond previous technological and economic limits.

## Data availability

All raw sequences used in this study have been deposited in the NCBI BioProject database with accession code PRJNA875933. Illumina 16S rRNA gene amplicon BioSamples: SAMN30633139-SAMN30633178. ONT 16S rRNA gene amplicon BioSamples: SAMN30633887-SAMN30633926. Illumina metagenomes were submitted to the BioSamples: SAMN30647025-SAMN30647033.

## Acknowledgements

The authors would like to thank the captain and crew of the R/V Atlantic Condor as well as Daniel Gibson and Greg Siddall from the Modular Ocean Research Infrastructure (MORI) project for a successful sampling expedition that made this work possible. Funding for ship time was provided by Nova Scotia Department of Natural Resources and Renewables, with support from Natural Resources Canada and Net Zero Atlantic. The work was supported by research funding from Genome Canadaʼs Genomics Applications Partnership Program facilitated by Genome Atlantic and Genome Alberta (to CRJH and AM), the Canada Foundation for Innovation (CFI-JELF 33752 to CRJH), and a Campus Alberta Innovates Program chair (CRJH). JZ was supported by post-doctoral fellowship awards from the Natural Sciences and Engineering Research Council (NSERC) and Mitacs. The authors also wish to thank Carey Ryan and Rhonda Clark for research and logistics support.

## CRediT Taxonomy Roles

Jackie Zorz: Formal Analysis, Conceptualization, Methodology, Software, Data Curation, Validation, Visualization, Writing – Original Draft Preparation

Carmen Li: Methodology, Investigation, Resources

Anirban Chakraborty: Investigation, Writing – Review and Editing

Daniel Gittins: Software, Validation, Writing – Review and Editing

Taylor Surcon: Investigation, Software, Validation

Natasha Morrison: Project Administration, Resources

Robbie Bennett: Project Administration, Supervision

Adam MacDonald: Funding Acquisition, Resources

Casey Hubert: Conceptualization, Funding Acquisition, Supervision, Writing – Original Draft Preparation

