## Supplementary Materials for "*SituSeq*: An offline protocol for rapid and remote Nanopore amplicon sequence analysis"

**S1 Table. Primer sequences used for ONT and Illumina PCR reactions.**

| Primer Set | Primer Name | 16S rRNA region | Sequence |
|---|---|---|---|
| **ONT** | 27F/1492R | V1-V9 | 5’ AGAGTTTGATCMTGGCTCAG 3’<br>5’ CGGTTACCTTGTTACGACTT 3’ |
| **Illumina** | 515F/806R | V4 | 5’ GTGYCAGCMGCCGCGGTAA 3’<br>5’ GGACTACNVGGGTWTCTAAT 3’ |

**S2 Table. Thermocycler programs for both ONT and Illumina PCR reactions.**

| Program | Temperature (°C) | Time (seconds) | Cycles |
|---|---|---|---|
| **ONT 16S** | 95 | 60 | 1 |
| | 95 | 30 | 25 |
| | 55 | 45 | |
| | 72 | 120 | |
| | 72 | 300 | 1 |
| **Illumina 16S** | 95 | 180 | 1 |
| | 95 | 30 | 10 |
| | 60 | 45 | |
| | 72 | 60 | |
| | 95 | 30 | 20 |
| | 55 | 45 | |
| | 72 | 60 | |
| | 72 | 300 | 1 |
| **Illumina Index** | 95 | 180 | 1 |
| | 95 | 30 | 10 |
| | 55 | 45 | |
| | 72 | 60 | |
| | 72 | 300 | 1 |

**S3 Table. Differentially abundant phyla identified for various groupings.** Calculated by the *multipatt* function in the package *indicspecies*. Significance codes: 0*** ; 0.001** ; 0.01*

| Test | Group | Phylum | Score |
|---|---|---|---|
| **ONT amplicon vs Illumina amplicon vs Illumina Metagenome (all sequences) (n=27)** | ONT amplicon | Dependentiae | 0.625** |
| | | Campylobacterota | 0.458* |
| | Illumina metagenome | Firmicutes | 0.739*** |
| | | RCP2-54 | 0.639*** |
| | | Patescibacteria | 0.602** |
| | | Synergistota | 0.597** |
| | | WPS-2 | 0.585* |
| | | Aquificota | 0.580* |
| | | Halanaerobiaeota | 0.511* |
| | | NKB15 | 0.475* |
| | | Deferrisomatota | 0.464* |
| | | Actinobacteriota | 0.459* |
| | | Poribacteria | 0.427** |
| | | AncK6 | 0.342* |
| | Illumina amplicon + Illumina metagenome | Calditrichota | 0.575** |
| | | Chloroflexi | 0.481* |
| | ONT amplicon + Illumina metagenome | Unknown | 0.734*** |
| | | Verrucomicrobiota | 0.492* |
| | | Margulisbacteria | 0.480* |
| **ONT amplicon vs Illumina amplicon (all sequences) (n=80)** | ONT amplicon | Unknown | 0.741*** |
| | | Dependentiae | 0.618*** |
| | | Margulisbacteria | 0.386*** |
| | | Verrucomicrobiota | 0.368*** |
| | | Campylobacterota | 0.333*** |
| | | Patescibacteria | 0.311** |
| | | MAT-CR-M4-B07 | 0.290** |
| | | Deferrisomatota | 0.285** |
| | | Modulibacteria | 0.244* |
| | Illumina amplicon | Chloroflexi | 0.438*** |
| | | Calditrichota | 0.362*** |
| | | Poribacteria | 0.329*** |
| | | NKB15 | 0.256** |
| | | Marinimicrobia (SAR406) | 0.241* |
| | | Proteobacteria | 0.232* |
| | | Halanaerobiaeota | 0.190*** |
| **Location – ONT samples (all sequences) (n=40)** | Purple Haze | WS2 | 0.770*** |
| | | Caldatribacteriota | 0.636** |
| | | Acetothermia | 0.586*** |
| | | Caldisericota | 0.535** |
| | | FCPU426 | 0.512* |
| | | Cloacimonadota | 0.511** |
| | | Desulfobacterota | 0.478* |
| | | Fermentibacterota | 0.421* |
| | | Desantisbacteria | 0.416* |
| | 175NW | Unknown | 0.907*** |
| | | Sumerlaeota | 0.875*** |
| | | Acidobacteriota | 0.849*** |
| | | Planctomycetota | 0.795*** |
| | | Armatimonadota | 0.679*** |
| | | Verrucomicrobiota | 0.672*** |
| | | Nitrospinota | 0.672*** |
| | | BH180-139 | 0.644** |
| | | Hydrogenedentes | 0.624*** |
| | | 10bav-F6 | 0.606** |

| | | | |
|---|---|---|---|
| | | GN01 | 0.596*** |
| | | WOR-1 | 0.587** |
| | | Aerophobota | 0.555** |
| | | Chloroflexi | 0.549** |
| | | Nitrospirota | 0.538** |
| | | Methylomirabilota | 0.532*** |
| | | NB1-j | 0.505* |
| | | Elusimicrobiota | 0.499** |
| | | Dadabacteria | 0.496** |
| | | SAR324 clade | 0.492*** |
| | | Gemmatimonadota | 0.490* |
| | | Marinimicrobia (SAR406) | 0.452* |
| | | AncK6 | 0.442*** |
| | | MBNT15 | 0.440* |
| | | Zixibacteria | 0.436* |
| | | Patescibacteria | 0.407* |
| | Tiny Bubbles + Clamshell | Actinobacteriota | 0.597** |
| | | Bacteroidota | 0.512** |
| | Kilo | Schekmanbacteria | 0.725*** |
| | | Deferrisomatota | 0.678*** |
| | | Modulibacteria | 0.665*** |
| | | Campylobacterota | 0.604** |
| | | DTB120 | 0.550** |
| **Location – Illumina samples (all sequences) (n = 40)** | Purple Haze | WS2 | 0.808*** |
| | | Acetothermia | 0.708*** |
| | | Cloacimonadota | 0.706*** |
| | | Caldatribacteriota | 0.681*** |
| | | CK-2C2-2 | 0.667*** |
| | | Dependentiae | 0.643*** |
| | | Fermentibacterota | 0.554** |
| | | Patescibacteria | 0.553** |
| | | Desantisbacteria | 0.521* |
| | | Caldisericota | 0.505* |
| | | Desulfobacterota | 0.486** |
| | | TA06 | 0.485* |
| | | FCPU426 | 0.478* |
| | | Marinimicrobia (SAR406) | 0.405* |
| | 175NW | Acidobacteriota | 0.917*** |
| | | Planctomycetota | 0.828*** |
| | | Armatimonadota | 0.670*** |
| | | Aerophobota | 0.664*** |
| | | Unknown | 0.643*** |
| | | Sumerlaeota | 0.616** |
| | | Methylomirabilota | 0.603*** |
| | | 10bav-F6 | 0.578** |
| | | BH180-139 | 0.563** |
| | | Nitrospinota | 0.561** |
| | | Nitrospirota | 0.559** |
| | | Chlorovlexi | 0.550** |
| | | Dadabacteria | 0.525** |
| | | WOR-1 | 0.495* |
| | | SAR324 clade | 0.493** |
| | | Hydrogenedentes | 0.492* |
| | | NB1-j | 0.492* |
| | | MBNT15 | 0.485* |
| | | Gemmatimonadota | 0.425* |
| | | Verrucomicrobiota | 0.397* |
| | | Actinobacteriota | 0.713*** |

| | | | |
|---|---|---|---|
| | Tiny Bubbles + Clamshell | Bacteroidota | 0.437* |
| | | Myxococcota | 0.406* |
| | Kilo | Deferrisomatota | 0.777*** |
| | | Campylobacterota | 0.689*** |
| | | DTB120 | 0.676*** |
| | | Poribacteria | 0.664*** |
| | | Fibrobacterota | 0.577** |
| | | NKB15 | 0.571** |
| | | Schekmanbacteria | 0.534** |
| | | Margulisbacteria | 0.431* |

**S4 Table. Differentially abundant phyla identified for various groupings.** Calculated by the *multipatt* function in the package *indicspecies* with the number of reads per sample subsampled to 1000. Significance codes: 0*** ; 0.001** ; 0.01*

| | | | |
|---|---|---|---|
| **Location – ONT samples (1000 sequences per sample) (n = 40)** | **Purple Haze** | **Caldatribacteriota** | **0.635*** ** |
| | | WS2 | 0.612*** |
| | | Acetothermia | 0.536** |
| | | Cloacimonadota | 0.485* |
| | | Desulfobacterota | 0.481* |
| | | Caldisericota | 0.450** |
| | 175NW | Unknown | 0.854*** |
| | | Sumerlaeota | 0.841*** |
| | | Acidobacteriota | 0.831*** |
| | | Planctomycetota | 0.810*** |
| | | Nitrospinota | 0.697*** |
| | | Armatimonadota | 0.642*** |
| | | Verrucomicrobiota | 0.615*** |
| | | BH180-139 | 0.588** |
| | | Hydrogenedentes | 0.568*** |
| | | Chloroflexi | 0.559** |
| | | 10bav-F6 | 0.550** |
| | | Elusimcrobiota | 0.525** |
| | | Methylomirabilota | 0.507*** |
| | | Nitrospirota | 0.496* |
| | | NB1-j | 0.493* |
| | | Gemmatimonadota | 0.493* |
| | | WOR-1 | 0.489** |
| | | Aerophobota | 0.480** |
| | | Bdellovibrionota | 0.476* |
| | | SAR324 clade | 0.475*** |
| | Tiny Bubbles + Clamshell | Actinobacteriota | 0.62*** |
| | | Bacteroidota | 0.54** |
| | Kilo | Schekmanbacteria | 0.705*** |
| | | Campylobacterota | 0.619** |
| | | DTB120 | 0.517* |
| | | Deferrisomatota | 0.438* |
| **Location – Illumina samples (1000 sequences per sample) (n = 40)** | Purple Haze | WS2 | 0.687*** |
| | | Cloacimonadota | 0.682*** |
| | | Caldatribacteriota | 0.682*** |
| | | Acetothermia | 0.673*** |
| | | Dependentiae | 0.557*** |
| | | CK-2C2-2 | 0.540** |
| | | Patescibacteria | 0.538** |
| | | Desulfobacterota | 0.472* |
| | | Fermentibacterota | 0.459* |
| | | TA06 | 0.452* |
| | 175NW | Acidobacteriota | 0.910*** |
| | | Planctomycetota | 0.811*** |
| | | Methylomirabilota | 0.599*** |
| | | Nitrospirota | 0.593** |
| | | Aerophobota | 0.583** |
| | | Armatimonadota | 0.583** |
| | | 10bav-F6 | 0.559** |
| | | Chloroflexi | 0.538** |
| | | Nitrospinota | 0.527** |
| | | Unknown | 0.515* |

| | | | |
|---|---|---|---|
| | | NB1-j | 0.492* |
| | | Dadabacteria | 0.468* |
| | | SAR324 clade | 0.449** |
| | | WOR-1 | 0.438* |
| | Tiny Bubbles + Clamshell | Actinobacteriota | 0.670*** |
| | | Myxococcota | 0.474* |
| | | Bacteroidota | 0.434* |
| | Kilo | Campylobacterota | 0.686*** |
| | | Fibrobacterota | 0.629** |
| | | Schekmanbacteria | 0.609** |

**Supplementary Data (separate files):**

S1 Data. *SituSeq* code

S2 Data. Sequences for species of interest BLAST search
